# Single Testicular Biopsy: Changes in the vascular architecture of the tunica albuginea

**DOI:** 10.1101/2024.02.26.582049

**Authors:** Antonio Barbosa de Oliveira Filho, Rita Luiza Peruquetti, Lucas Trevizani Rasmussen, Marcos Teixeira Cesar, Crescêncio Alberto Pereira Cêntola, Agnaldo Pereira Cedenho

## Abstract

Open testicular biopsy is a commonly indicated procedure for diagnosis of testicular histopathology prior to referral to testicular sperm extraction (TESE) or microsurgical testicular sperm extraction (M-TESE) in men with non-obstructive azoospermia (NOA). There are known hematological changes resulting from the rupture of the blood–testis barrier in the TESE and M-TESE procedures, but changes to the microvasculature have not yet been described when the open biopsy is performed prior to spermatozoid collection for intracytoplasmic sperm injection. In this experimental study, open biopsy was performed on testes from adult rabbits. After a 45 days recovery period, the animals underwent a laparotomy for the surgical removal of the aortic patch, both gonadal arteries, the vasa deferentia, and both testicles. The laparotomy was followed by an angiography. The comparative results demonstrate a breakdown in the vascularization of the testicles biopsied relative to the microvasculature pattern of the tunicae albugineae of the intact testicles. The vascular damage resulting from open testicular biopsy reflects one more negative impact on spermatogenic function of previously lesioned testicles besides the known alterations (fibrosis, decrease in testosterone levels, hematological changes). Therefore, we suggest that the M-TESE should be the choice method for sperm retrieval in men with NOA and it should be planned and performed with simultaneous diagnostic and therapeutic objectives in order to increase the patient’s chances of reproductive success.

## Introduction

Individuals with non-obstructive azoospermia (NOA) are diagnosed based on ejaculated semen obtained via masturbation when lack of spermatozoa is detected. The assessment of these individuals also considers the amount of gonadotropins (FSH and LH) as a non-invasive predictive criterion when investigating the possible existence of remaining evidence of spermatogenesis (Jarow et al., 1989).

Diagnostic unilateral or bilateral open testicular biopsies in patients with NOA are often requested after gonadotropin and total testosterone levels are determined. The objectives of the diagnostic biopsy are to predict the existence of any foci of spermatogenesis and to provide an assessment of the anatomical pattern responsible for the azoospermia (Glina et al., 2005). The main anatomical patterns found in the biopsies of testicles from NOA patients are: (a) hypospermatogenesis; (b) maturation arrest; and (c) Sertoli cell-only (SCO) syndrome (Schulze et al., 1999). However, the anatomical pattern cannot be generalized to definitively label patients as having NOA or as being incapable of exhibiting spermatogenesis in regions not covered by the biopsy (Dabaja and Schlegel, 2013).

In 1993, the testicular sperm extraction (TESE) technique was developed for obtaining spermatozoa from open testicular biopsies performed on an experimental basis (Schoysman et al., 1993). The classic description of TESE states that three 1.0 cm transverse incisions must be made on the anterior side of the testicle (Silber et al., 1997; Silber et al., 1995). Despite the knowledge on the damages that this technique can cause as a result of a broken blood–testis barrier (BTB), TESE is routinely performed. A study that evaluated the number of TESE interventions necessary to obtain viable spermatozoa from men with NOA found that an average of three procedures are required to reach a sufficient rate of viable spermatozoid retrieval for intracytoplasmic sperm injection (ICSI), a finding which provides evidence of increased damage to an already compromised organ (Dadkhah et al., 2013). In an attempt to create new criteria for detecting potential chances of success through the use of TESE, other factors able to predict viable spermatozoid retrieval for ICSI have been evaluated. These include the analysis of total testicular volume and the levels of both FSH and inhibin B (Boitrelle et al., 2011).

In 1999, a microsurgical testicular sperm extraction (M-TESE) was proposed in order to improve upon the TESE technique by increasing the rate of viable spermatozoid retrieval per procedure, thus providing a higher likelihood of success with a single procedure (Schlegel, 1999). The M-TESE procedure consists of a surgical approach using a surgical microscope, which enables the opening of the tunica albuginea with magnification and the exploration of a larger area of the testicular parenchyma. In this method, the typical clusters of seminiferous tubules, which are markers of foci of spermatogenesis, can be visualized and collected. In addition to the wider exploration of the parenchyma under a microscope, another advantage of M-TESE over TESE is the reduction in damage to tunica albuginea vascularization (Dabaja and Schlegel, 2013; Franco et al., 2016; Okada et al., 2002).

A 2012 study reported that, in patients with severe NOA and histopathological results reflecting low chances of viable spermatozoid retrieval, the combination of conventional TESE (with three testicular incision points) and M-TESE produced a 66.2% rate of retrieval of viable spermatozoa for use in ICSI, which increases the chance of success in these cases (Marconi et al., 2012).

There have been substantial improvements in the techniques used for obtaining spermatozoa from NOA patients for use in ICSI. These advances provide greater chances of spermatozoid retrieval with less risk of damage during the procedure (Franco et al., 2016). These descriptions of the procedures used to obtain spermatozoa reflect the fact that all of the techniques break the BTB, which may have possible indirect repercussions for the immune system and which may cause direct damage to both tunica albuginea vascularization and the testicular parenchyma.

Therefore, the aim of the present study was to describe the testicular microvascular changes caused by the open testicular biopsy usually performed prior to TESE or M-TESE in men with NOA. Our results may point to any additional damages caused by the surgical intervention to the patient’s testicle, which may reduce any remaining foci of spermatogenesis decreasing their reproductive success chances.

## Material and Methods

The experimental procedures were carried out on twelve adult male New Zealand rabbits (±7.5 months / ±3,100g). During the experiments specimens were housed under standard conventional conditions (25°C; 40–70% relative humidity; 12 h light ⁄ 12 h dark). They were also monitored for signs of distress, pain, and/or infection on a daily basis, and they were given ad libitum access to food and water. Cages were cleaned on a weekly basis and/or when visibly soiled in order to maintain a clean environment. All of the procedures used for euthanasia were consistent with the Brazilian Guidelines for the Care and Use of Animals for Scientific and Educational Means (CONCEA, 2013). This study was approved by the Institutional Review Board from the School of Medicine of São José do Rio Preto (FAMERP).

Animals were separated into four groups (n=3/group): (1) Control group (CG): open testicular biopsies on the left testicle; (2) Sham group (SG): three small incisions on the left testicle followed by open testicular biopsies on the right testicle; (3) Experimental group (EG): M-TESE on the right testicle followed by open testicular biopsies on the left testicle; (4) Vascular control group (VCG): both testicles intact. All the groups were acclimatized for ten days at the animal facility before surgical procedures. After a 45-day recovery period, animals from EG group as well as animals from VCG underwent laparotomy and bilateral orchiectomy for the angiographic study of both testicular arteries and of the anatomical pattern of the tunica albuginea.

### Surgical Procedures

#### Testicular Biopsy (EG)

Experimental procedure was performed at the vivarium of the School of Medicine of São José do Rio Preto, São José do Rio Preto, São Paulo, Brazil, after the animals were fasted for 12 hours. The animals in the experimental group (EG) were anaesthetized through intramuscular administration of ketamine hydrochloride and xylazine (1.0 to 1.5 ml/animal). After the success of the anesthetic was confirmed, each animal received a trichotomy, antisepsis, surgical incision, and testicle exteriorization. The area around the surgical field was covered with a sterile silicone towel. A small vertical incision (≅1.0 cm) was made through the scrotal skin and through the tunica albuginea using a no. 11 blade, and two small pieces of the left testicle tissue were removed. Sizes of the surgical incision were 0.5 larger then surgical incision usually performed on human open testicular biopsy. Testicular fragments from both testicles of animals in all the groups were fixed in Bouin’s fixative solution for further study. The tunica albuginea and the skin of the scrotum region were sutured using Prolene® no. 7-0 sutures (Figure 1). After recovering from the surgical procedure, the animals were taken to their cages and were housed under standard conventional conditions for 45 days. After this period, the animals underwent the procedures described below.

**Figure 1.**
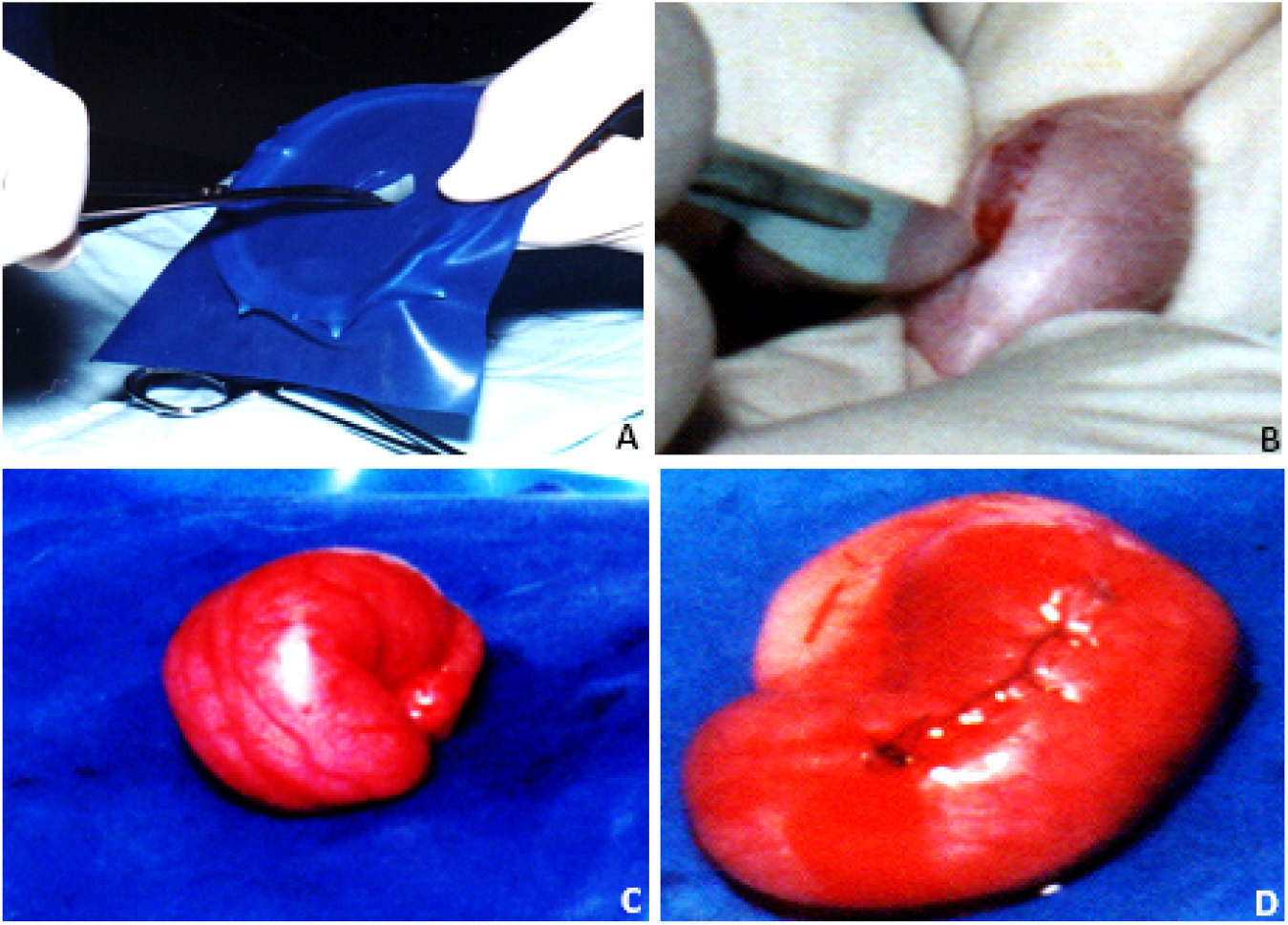
Surgical procedure to perform single incision open testicular biopsy. (A) The area around the surgical field covered with a sterile silicone towel. (B) A small surgical cut through the scrotal skin and through the tunica albuginea for exposure of the left testicle. (C) Left testicle exposed and ready to be biopsied. (D) The tunica albuginea and the skin of the scrotum region after suturing procedure.

### Laparotomy and Bilateral Orchiectomy (VCG and EG)

The same preoperative procedures used for testicular biopsy were performed. The posterior peritoneum was sectioned using a surgical microscopy. Next, the large blood vessels (the aorta and the vena cava) and the right and left gonadal arteries were exposed and dissected. These vessels were removed with the testicles, and these biological materials were kept at 5°C in a physiological solution. Animals were euthanized through respiratory arrest using a high anesthetic dosage after the surgical procedures.

### Angiography of the Testicular Artery and Tunica Albuginea (VCG and EG)

Samples collected after laparotomy and bilateral orchiectomy were taken to the Radiology and Angiography Facility (CRIVA) of Beneficiência Portuguesa Hospital in São José do Rio Preto, São Paulo, Brazil. Angiography was performed on the testicular arteries and tunica albuginea through the use of a no. 5 polyethylene catheter and a cotton line in both extremities (inferior and superior) of the aortic patch. Contrast solution consisting of BARIOGEL® 100% barium sulfate solution, and India ink was injected using a 5 ml syringe. Contrast injection was followed by radioscopy, and images were documented on radiography film (Figure 2).

**Figure 2.**
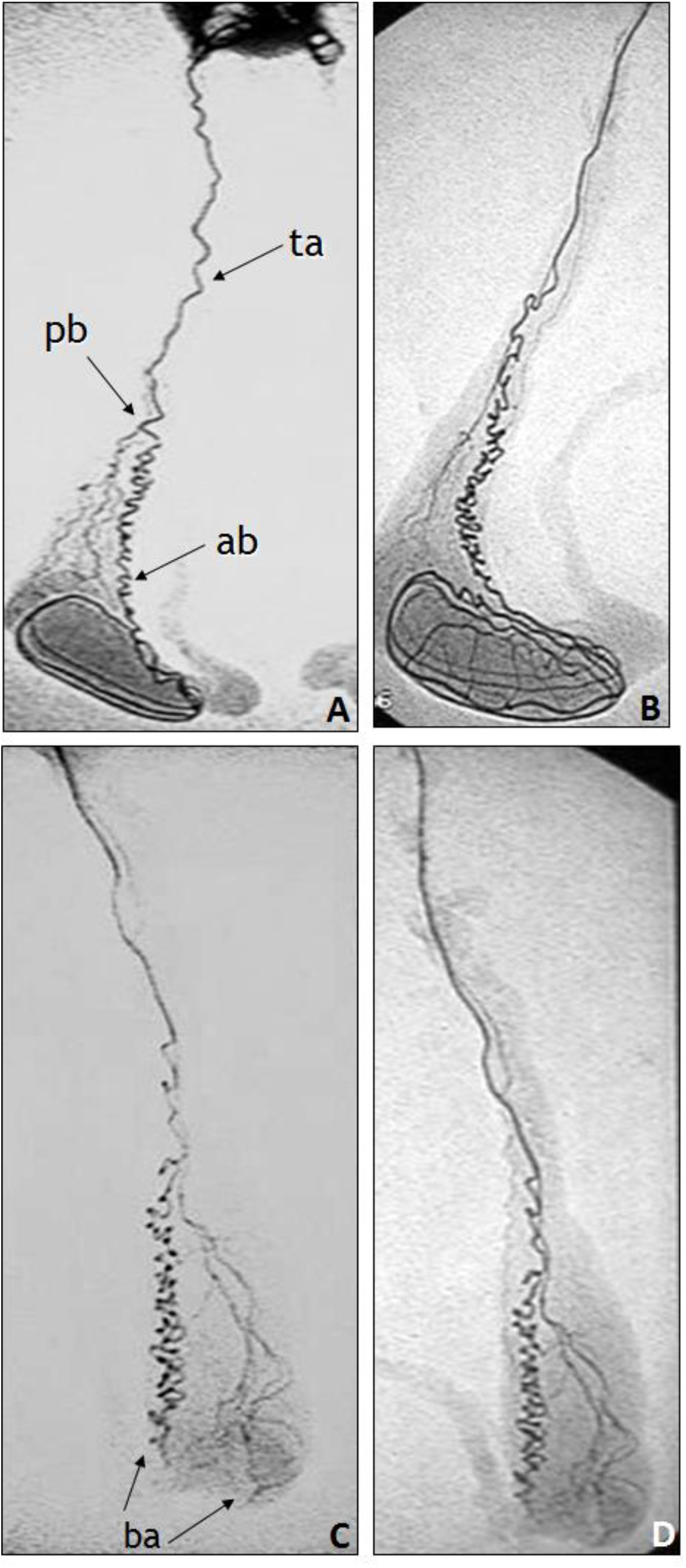
Angiography of the testicular and epididymal arteries as well as the tunica albuginea of the right intact testis - VCG (A and B) and left testis after a 45 day recovery from single open biopsy - EG (C and D). (A) and (C) represent radiographs obtained at the beginning of contrast injection and (B) and (D) represent radiographs obtained after the completion of contrast injection. It is possible to observe the pre-testicular and testicular angiographic pattern in the intact testicles (A and B). In the testicles biopsied (C and D), it is possible to notice that angiography prior to the biopsy site (ba) is preserved, but there is a compromise in the blood flow of the tunica albuginea in the regions that follows the incision site. (ta) = testicular artery; (pb) = posterior branch; (ab) = anterior branch; (ba) = biopsy area.

## Results

### Control Group (VCG)

Vascularization of the testis was clear from the origin of the gonadal artery coming from the abdominal aorta and through its path along the vas deferens (pre-testicular segment) and also along its entire route in the tunica albuginea and its sub-capsular anastomoses (Figures 2A and 2B). In the pre-testicular segment, the gonadal artery was found to fork at the distal third of its path, causing posterior branch flow and forming the subsequent epididymal artery. At this point, the gonadal artery becomes highly curved and coiled (Figures 2A and 2B).

The epididymal artery protrudes on the medial portion of the testicle through the cranial to caudal region of the testis and becomes less coiled as it forms three circuits in the longitudinal direction of the testis (Figures 2A and 2B). After covering the three circuits in the longitudinal direction of the testis, the blood flows through small vessels that come together at joining the circuit in the transverse direction producing an angioarchitecture that resembles a mesh. The posterior epididymal artery is smaller in diameter and originates from the distal third of the gonadal artery, running parallel to the gonadal artery in its distal path, and projecting into the caudal portion of the epididymis. The posterior epididymal artery is also smaller in diameter than the gonadal artery, and, like the gonadal artery, it arises from the aorta artery and projects itself onto the anterior portion of the epididymis.

### Experimental Group (EG)

There was no difference in the vascularization patterns of the gonadal and epididymal arteries in the pre-testicular segment 45 days after the open testicular biopsy (Figures 2C and 2D). However, there was an interruption in two of the testicles’ longitudinal circuits, and some branches disappeared (Figures 2C and 2D). The presence of only one circuit without interrupted blood flow and a no mesh-like appearance was found to be different from the angioarchitecture of the intact testicles (Figures 2C and 2D). The posterior and anterior epididymal arteries were identified and preserved and exhibited integrity of the blood supply to the epididymis in their cranial and caudal portions.

### Testicular Histology and Morphometry (CG, SG, EG, VCG)

There was an increase of lower Johnsen-like scores (6 and 5) in the seminiferous tubules from left testicle from EG indicating that the surgical procedure has caused spermatogenic alterations, leading to maturation arrest in this group. Morphometric evaluation of the seminiferous tubules did not demonstrate important differences among the groups. These results were already published in a previous paper (see Oliveira et al. (2010) for more details).

## Discussion

The objective of this study was to describe changes in testicular and tunica albuginea vascularization after open testicular biopsy in order to predict its clinical impact for men with NOA who are candidates for ICSI.

Open testicular biopsy for diagnostic purposes is a procedure routinely employed by many professionals, despite the fact that this procedure does not provide convincing evidence of foci of spermatogenesis in any portion of the testicular parenchyma through the collection of a small sample from the seminiferous tubules (Dabaja and Schlegel, 2013). Bettocchi et al. (1998) concluded that biopsies revealing patterns that corresponded to bilateral Sertoli cell-only syndrome does not guarantee the absence of spermatogenesis, and also that FSH levels are not solely good indicator of spermatogenesis progression. However, the effects of these open biopsies on the testicles are often disregarded due to a lack of knowledge of its effect on the anatomy of the tunica albuginea microvasculature (Franco et al., 2016).

Schlegel and Su (1997), found that after TESE, whose incision is very similar to the incision used in open testicular biopsy, the lesioned area presents hematomas, fibrosis, and a loss of vascularization. Evidence of these abnormalities can be seen up to six months after surgery. Most of the pathophysiological responses associated with fibrotic damage similar to that observed in the testicle after open testicular biopsy are caused through the action of mast cells, which are multifunctional cells in the immune system, whose physiological imbalance may compromise the blood testicular barrier (BTB) function (Haidl et al., 2011). The BTB essentially consists of tight junctions between the adjacent Sertoli cells that keeps germ cells in more advanced stages within the testicle and protects them from immune cells, which may recognize germ cells as foreign bodies due to their genetic differences. This barrier breaks down in response to stimulation from FSH and androgens to allow for the passage of mature cells to luminal regions of the seminiferous tubules, and its function is essential to the effective completion of the spermatogenesis process (Stanton, 2016).

Changes in testicular microvasculature demonstrated in this study indicates that open testicular biopsy may damage the BTB and, in doing so, may negatively affect the possible foci of spermatogenesis remaining in the testicle in two distinct ways: by increasing mast cell actions in the postoperative fibrosis region, which may affect the physiology of the BTB, and through vascular changes affecting FSH and androgen levels, as these hormones play a fundamental role in regulating the BTB and enabling the movement of advanced-stage germ cells to the luminal region of the seminiferous tubules. Thus, it is clear that any remaining foci of spermatogenesis will be compromised, further contributing to infertility. It’s important to be aware of some limitations of this study, such as: (1) differences on the size of surgical incision performed in our open biopsy when compared to human procedure; (2) Larger testis weigh/body ratio in rabbits when compared to human; (3) Short follow-up period period after testis retrieval (Fedder et al., 2017); (4) Absence of angiography for the sham (SG) and for the control groups (CG).

There is an apparent consensus in the literature that the M-TESE procedure is an excellent choice in cases of NOA—even in cases of Klinefelter syndrome, cryptorchidism, Y chromosome AZFc microdeletion (which may require preoperative hormone treatment), or cases involving chemotherapy. Immediate M-TESE increases the chances of obtaining viable spermatozoa for processing and storage for future ICSI (Dabaja and Schlegel, 2013; Franco et al., 2016; Kalsi et al., 2011). Though M-TESE is a relatively expensive and complicated technique involving the microscopic identification of the regions of interest, this technique increases the chances of viable spermatozoid collection, reduces damages to testicular tissue, and may be used to collect samples for a histopathological diagnosis with simultaneous spermatozoid retrieval.

For professionals who prefers to obtain the histopathological diagnosis before indicating a spermatozoid collection procedure, one option could be to perform a diagnostic biopsy of testicle fragments obtained using the testicular fine-needle aspiration (TESA) technique (Lewin et al., 1999). Some articles in the literature have reported that, while TESA is a simple and inexpensive method with fewer surgical complications than TESE, it should be avoided because a lower percentage of viable spermatozoa are retrieved from men with NOA. However, when other factors are considered, such as patients’ FSH levels (<15 IU/I) and/or testicular pathology of hypospermatogenesis, TESA has been found to be as effective as TESE in terms of viable spermatozoid retrieval (Nowroozi et al., 2012).

Furthermore, the development of an artificial neural system and a nomogram based on clinical data from potential spermatozoid retrieval patients, such as age, FSH levels, testicular volume, and history of cryptorchidism, Klinefelter syndrome, and/or varicocele, has resulted in a 59% success rate in the prediction of viable spermatozoid retrieval. This system is therefore a good, non-invasive tool for selecting patients for spermatozoid collection without damaging the already unstable organ (Ramasamy, 2013).

To summarize, open testicular biopsy performed in our experimental group (EG) produced disorganization of testicular microvasculature. Interruption in the communication between the testicular artery and its posterior branch was also observed as well as a substantial reduction in the caliber of the other branches of the testicular parenchyma. These findings show that testicular biopsy is not a risk-free procedure. We would therefore like to propose different conduct for patients with NOA who are candidates for TESE or M-TESE to avoid additional damage and potential negative effects on testicles with remaining foci of spermatogenesis. Based on our findings, we propose that M-TESE should be used as a mixed approach that is both therapeutic and diagnostic for NOA patients.

## Acknowledgments

We would like to thank Dr. Tiago da Silveira Vasconcelos and Danielle Jacqueline Deremo Cosimo for the critical reading of the manuscript. We would also like to thank all of the members of Dr. Cedenhos’s Laboratory (UNIFESP) for their help, particularly with regents and in discussing the research.

